# Direct nanopore sequencing of mRNA reveals landscape of transcript isoforms in apicomplexan parasites

**DOI:** 10.1101/2020.02.16.946699

**Authors:** V Vern Lee, Louise M. Judd, Aaron R. Jex, Kathryn E. Holt, Christopher J. Tonkin, Stuart A. Ralph.

**Affiliations:** Department of Biochemistry and Molecular Biology, Bio21 Molecular Science and Biotechnology Institute, The University of Melbourne, Melbourne, Victoria, 3010, Australia; The Walter and Eliza Hall Institute of Medical Research, Parkville, Melbourne, Victoria 3052, Australia.; Department of Infectious Diseases, Central Clinical School, Monash University, Melbourne, Victoria, Australia; Faculty of Veterinary and Agricultural Sciences, The University of Melbourne, Parkville, Victoria, Australia; London School of Hygiene and Tropical Medicine, London WC1E 7HT, UK; Department of Medical Biology, The University of Melbourne, Melbourne, Victoria 3052, Australia.

## Abstract

Alternative splicing is a widespread phenomenon in metazoans by which single genes are able to produce multiple isoforms of the gene product. However, this has been poorly characterised in apicomplexans, a major phylum of some of the most important global parasites. Efforts have been hampered by atypical transcriptomic features, such as the high AT content of Plasmodium RNA, but also the limitations of short read sequencing in deciphering complex splicing events. In this study, we utilised the long read direct RNA sequencing platform developed by Oxford Nanopore Technologies (ONT) to survey the alternative splicing landscape of *Toxoplasma gondii* and *Plasmodium falciparum*. We find that while native RNA sequencing has a reduced throughput, it allows us to obtain full-length or near full-length transcripts with comparable quantification to Illumina sequencing. By comparing this data with available gene models, we find widespread alternative splicing, particular intron retention, in these parasites. Most of these transcripts contain premature stop codons, suggesting that in these parasites, alternative splicing represents a pathway to transcriptomic diversity, rather than expanding proteomic diversity. Moreover, alternative splicing rates are comparable between parasites, suggesting a shared splicing machinery, despite notable transcriptomic differences between the parasites. This work highlights a strategy in using long read sequencing to understand splicing events at the whole transcript level, and has implications in future interpretation of RNA-seq studies.

## Introduction

Transcriptomic analyses have been central to insights into the biology and pathogenesis of eukaryotic pathogens. The best-characterised eukaryotic pathogen transcriptomes are those of the phylum Apicomplexa. This phylum includes some of the most important parasites impacting human and veterinary health, such as *Plasmodium* and *Toxoplasma*. *Plasmodium* is the causative agent of malaria, a devastating parasitic disease infecting over 200 million individuals and killing 400,000 each year [1]. *Toxoplasma* causes toxoplasmosis, a widespread zoonoses that primarily impacts immunocompromised, young, and pregnant individuals [2], and is thought to infect a third of the world’s population [3]. The pathogenesis of apicomplexan infections is intimately linked to the parasites’ life cycles. The life cycle of most parasitic apicomplexans is complex, involving multiple differentiated forms and hosts, and this requires reprograming of the parasite transcriptome.

Early transcriptomic experiments sought to utilise techniques such as microarrays and Sanger sequencing of cDNA or EST libraries to understand changes in gene expression that define the pathogenesis of the parasites. These studies reveal that the timing of appearance and abundance of individual mRNAs follow developmentally distinct patterns [4], even for many predicted housekeeping genes. For example, the expression of the actin gene family in *P. falciparum* is developmentally tuned, with actin I primarily transcribed in asexual intraerythrocytic life stages while actin II is primarily present in sexual stage parasites [5, 6]. Unusually, however, there is a poor correlation between protein and mRNA expression profiles for many genes in parasitic apicomplexans [7]. In one experiment, Foth and colleagues found widespread discrepancies between temporal expression patterns of proteins and transcripts in *P. falciparum* [8]. Such discrepancies suggest that substantial post-transcriptional regulation occurs within these parasites. Indeed, with the advent of RNA-seq, more recent studies now show that multiple layers of gene expression are required for parasite life progression, through transcriptional, post-transcriptional, and epigenetic control mechanisms [9–11].

RNA splicing provides one such source of co- and post-transcriptional regulation. In this process, introns are removed from the pre-mRNA and the exons retained to form one contiguous molecule that is then translated by the ribosome. However, for complex mRNAs, alternative splicing either of untranslated regions, or the exonic chain, can add additional complexity. Through this process, pre-mRNA species can be differentially spliced, to create multiple distinct mature mRNAs from a single gene. This can alter regulation of the gene, for example by removing small-RNA binding sites [12] or diversify the proteome, as individual genes may encode multiple protein isoforms with altered structure or function [13].Indeed, proteomic analyses have revealed widespread protein isoforms arising from single genes, corresponding with varying activity, stability, localisation and post-translational modifications [14, 15]. With advances in genome and transcript sequencing, it has become apparent over the last decade that alternative splicing of pre-mRNA occurs to a great extent. For example, more than 95% of human genes are alternatively spliced, and many transcript isoforms are specific to tissues or cellular states [16]. Such observations suggest that RNA diversity is more complex than previously appreciated [17].

Although alternative splicing appears to play a major (though debated) role in post-transcriptional control in metazoans, the process in less understood in apicomplexans. Studies have identified apicomplexan genes with crucial alternative splicing outcomes [18]. For example, alternative splicing is required for attaching a protein trafficking pre-sequence onto two adjacent gene coding sequences [19], and normal multi-organellar targeting of the *P. falciparum* cysteinyl tRNA synthetase, which is essential for parasite survival [20]. Nonetheless, there is little other study of alternative splicing in this phylum. Understanding diversity of parasites transcripts is crucial for drug and vaccine development because certain putative target genes may produce isoforms that escape the intervention. This has been postulated for the chloroquine resistance transporter gene of *P. falciparum* (*Pf*CRT) in clinical isolates, though the role of the splice variants remains unclear [21]. In other organisms, there is some evidence showing that essential genes are more likely to have alternatively spliced transcripts compared to non-essential genes [22, 23]. This has not been explored in apicomplexans but highlights further considerations for investigating drug targets and interventions.

The lack of data for apicomplexan gene isoforms is a major obstacle to dissecting the complexity of transcript outcomes. Traditionally, transcriptomic studies employing RNA-seq have relied on short read technologies such as Illumina, 454 and Ion-torrent [24]. Despite the power of very high sequencing depth and low error rates, the short reads present a limitation in that simultaneously occurring alternative-splicing events within individual transcripts cannot be unambiguously detected or linked. Previously-developed computational methods for full-length transcript assembly from short read sequencing data are often computationally intensive, and can produce ambiguous or conflicting results between different algorithms [25]. In addition, sequencing on cDNA strands amplified by PCR has a propensity to introduce biases in relative transcript abundances and rare isoform identification [26]. Hence, it is difficult to draw functional relationships between simultaneous alternative splicing events and observable phenotypes. In apicomplexan parasites, simultaneously occurring alternative splicing events within a specific transcript isoform do occur [27]. However, the studies that unearthed these transcript isoforms relied on cDNA probes and reverse transcription PCR, and the wider extent of this phenomenon is unknown.

Recently-developed third generation sequencing platforms, such as those developed by Oxford Nanopore Technologies (ONT) and Pacific Biosciences (PacBio), are capable of producing significantly longer reads at the single-molecule level. These technologies have been used in various applications such as resolving genomic and transcriptional landscapes [28, 29], single cell transcriptome sequencing [30], and DNA or RNA methylation pattern profiling [31, 32]. PacBio has recently been used to generate an amplification-free transcriptome from *Plasmodium falciparum* cDNA, which has helped to elucidate transcriptional start sites and to improve annotation of the 5’and 3’ UTRs [33]. Unlike most other sequencing platforms, a notable characteristic of ONT sequencing is the ability to directly sequence native RNA [34]. With this methodology, each read represents a complete molecular transcript, which could thus significantly resolve weaknesses of amplification-based RNA-seq. In particular, each spliced isoform need only be counted as individual reads, as opposed to complex assignment and assembly of multiple spliced reads. Furthermore, due to differences between DNA and RNA molecules, contaminating DNA sequences cannot be correctly base-called after sequencing and so are easily discarded [35]. Recently, several studies have successfully applied single molecule, long-read sequencing to identify a high number of novel transcript isoforms [28, 36, 37]. However, these studies have also identified several caveats including a reduced throughput and high error rates.

In this study, we evaluate the ability ONT direct RNA sequencing to characterise the alternative splicing landscape of two parasitic apicomplexans, *T. gondii* and *P. falciparum.* Our analyses show that alternative splicing, particularly intron retention, is extensive throughout the transcriptome, with most multi-exon genes having some degree of intron retention, and some genes only rarely producing transcripts with all introns removed. The long reads produced from ONT sequencing showed that most of these alternative splicing events are likely non-productive in protein-coding capacity, but may provide an additional layer of gene expression regulation.

## Results

### Direct RNA sequencing of *T. gondii* and *P. falciparum* allows the detection of full-length transcripts

We generated ONT sequencing reads of polyA-selected RNA from asynchronous *T. gondii* (Pru) tachyzoites and *P. falciparum* (3D7) mixed asexual intraerythrocytic stage parasites. Mixed-stage cultures were used to maximise transcript diversity for these samples. For *T. gondii*, we obtained a total of 310,813 reads corresponding to about 500 million bases (Mb). For *P. falciparum*, we obtained a total of 456,098 reads, corresponding to about 300 Mb of data. Although the *P. falciparum* sample yielded fewer sequenced bases, we estimate the theoretical gene coverage for both parasite samples to be similar at 25-26 fold due to differences in gene number and length. Using minimap2 [38], we successfully mapped 78.90% of the *T. gondii* reads, and 44.48% of the *P. falciparum* reads to the parasite genomes. We analysed the quality of the sequencing reads using FASTQC, and found consistently high-quality scores over the length of reads, with no drop-off in quality even at reads of 10 Kb (Fig 1B). This is important because base quality scores generally correlate with read accuracy to the reference sequence. That is indeed the case for our dataset (supplementary figure S1). We used AlignQC to estimate the base-call error rate of the transcript reads based on aligned segments and found, on average, an error rate of 19-20% for both parasites. The read-length distributions of the mapped and unmapped data (Fig 1A) show that the mapped reads are predominantly longer than the unmapped reads, with some read lengths exceeding 10 kb. As expected, there was a sharp increase in unmapped read counts at the ~1.35 kb length (Fig 1A) corresponding to yeast enolase 2, a calibration standard added during the library preparation. We calculated the average mapped read lengths to be ~1.9 k and 1.3k bases for *T. gondii* and *P. falciparum* respectively, well within previous range estimates of predicted transcript lengths of both organisms [39].

**Figure 1.**
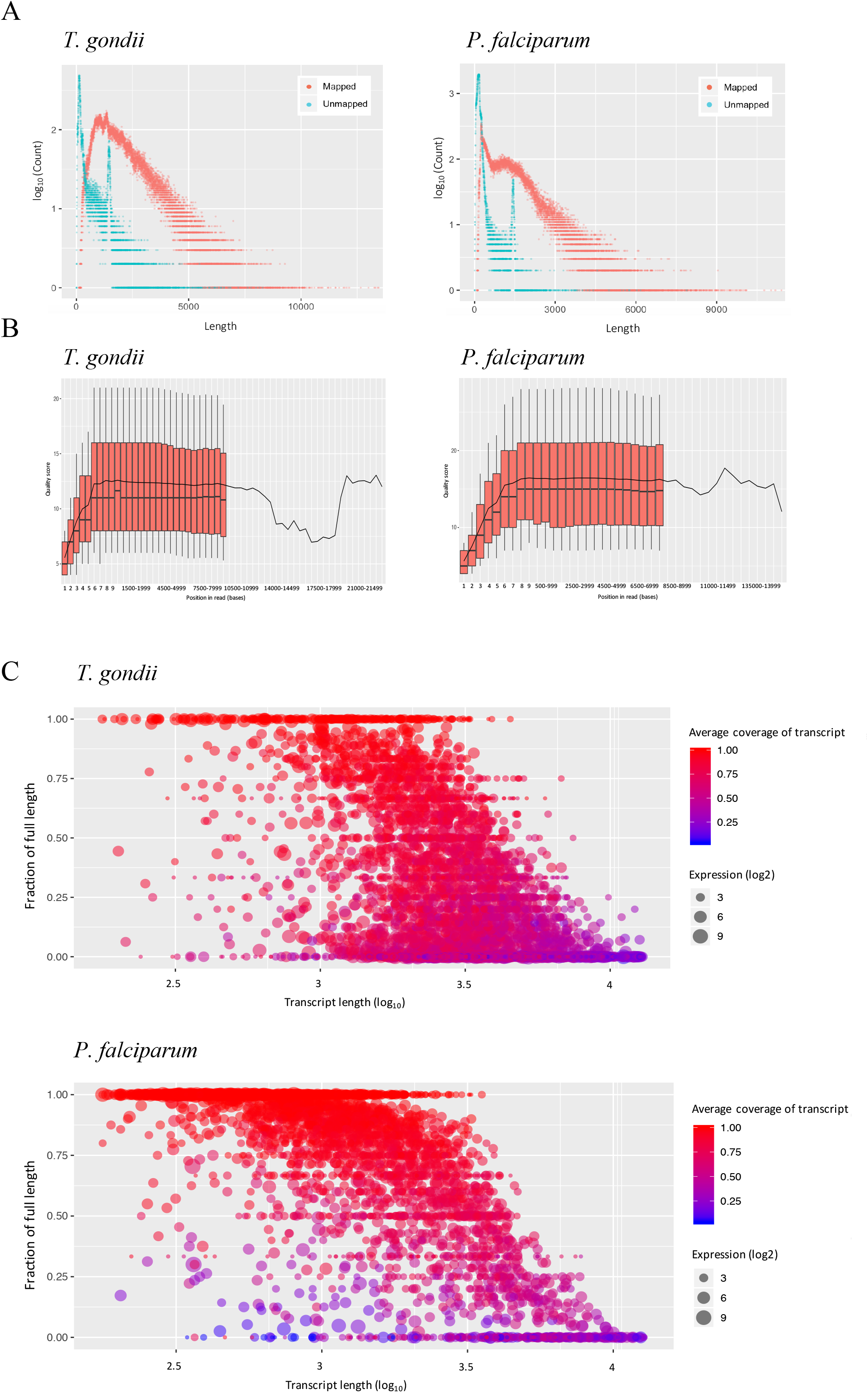
Summary of the ONT direct RNA sequencing data from *T. gondii* and *P. falciparum*. Scatterplot of read length distribution of mapped (red) and unmapped (blue) reads. (B) Boxplot of quality scores across all bases at each position of the mapped sequencing reads. (C) Bubble scatter heat plots of the fraction of full length transcripts against transcript length. Size and color denotes expression and average coverage respectively.

To evaluate the capability of ONT sequencing to generate full-length transcript reads, we re-mapped the reads to the parasites’ annotated transcriptome and calculated the fraction of full-length transcripts per gene. We define full-length transcripts as reads that cover more than 95% of the predicted canonical transcript based on the annotation file. As illustrated in the Figure 1C, many transcripts were observed to have full-length reads, more so for the *P. falciparum* data. In *T. gondii*, 1117 genes have 75% or more of their corresponding reads that were considered full length. In *P. falciparum,* this number is 1835 genes. The difference can be attributed to the absence of 5’ and 3’ UTR annotations for *P. falciparum*, which results in the underestimation of predicted transcript lengths. In both cases, the fraction of mapped reads corresponding to full-length transcripts fell with increasing transcript length, independent of expression levels, which is consistent with read truncation disproportionately affecting the longest reads. To better understand the overall distribution of transcript coverage, we calculated the average coverage of the transcripts per gene. Again, there is a general decrease in transcript coverage with increasing transcript length (Figure 1C). However, many of the genes retain a high level of transcript coverage, even when there is a low fraction of full length reads. For example, in *T. gondii*, genes with predicted transcript lengths of 3 kb or longer only had an estimated 12% of their reads that were considered full length on average, even though the average read coverage for those genes is 50.64%. The overall average transcript coverage is calculated to be over 60% in both parasites (*T. gondii*: 64.73%, *P. falciparum*: 80.51%). Together, the data indicate a generally high proportion of full- or near full-length transcript reads.

### ONT sequencing is comparable with traditional RNA-seq for quantifying gene expression levels

To investigate the utility of ONT data to measure transcript abundance, we computed read-count correlations between our ONT dataset and reanalysed published Illumina-based RNA-seq datasets on comparable parasite samples. In theory, ONT RNA sequencing reads directly correspond to complete transcripts and so quantifying the expression of genes can be done by simple counting of the assigned reads. This is dissimilar to traditional short read RNA-Seq which necessitates a further normalisation step (e.g., reads or fragments *per kilobase of transcript* or *transcripts per million*) to account for the higher number of reads that would be generated from longer transcripts. For *T. gondii*, we used Illumina datasets from a closely related strain (ME49) due to a relative lack of comprehensive datasets for the Pru strain. Strikingly, we observe strong positive correlations between the ONT and Illumina datasets (Fig 2A & B), regardless of mapping to the transcriptome or genome (Spearman’s rho = 0.81 for transcriptome, 0.87 for genome). For *P. falciparum* we correlated the mixed stage ONT dataset with individual datasets from three main developmental stages (rings, trophozoites and schizonts), and a final combined dataset. In all cases, we found moderately positive correlations between the datasets. As expected, the correlation is higher in the later stages (supplementary figure S2; Spearman’s rho = 0.47 rings, 0.57 for trophozoites, 0.63 for schizonts), and the highest with the combined dataset (Figure 2A/B; Spearman’s rho = 0.64 for transcriptome, 0.68 for genome), a reflection of mRNA abundance in these different stages.

**Figure 2.**
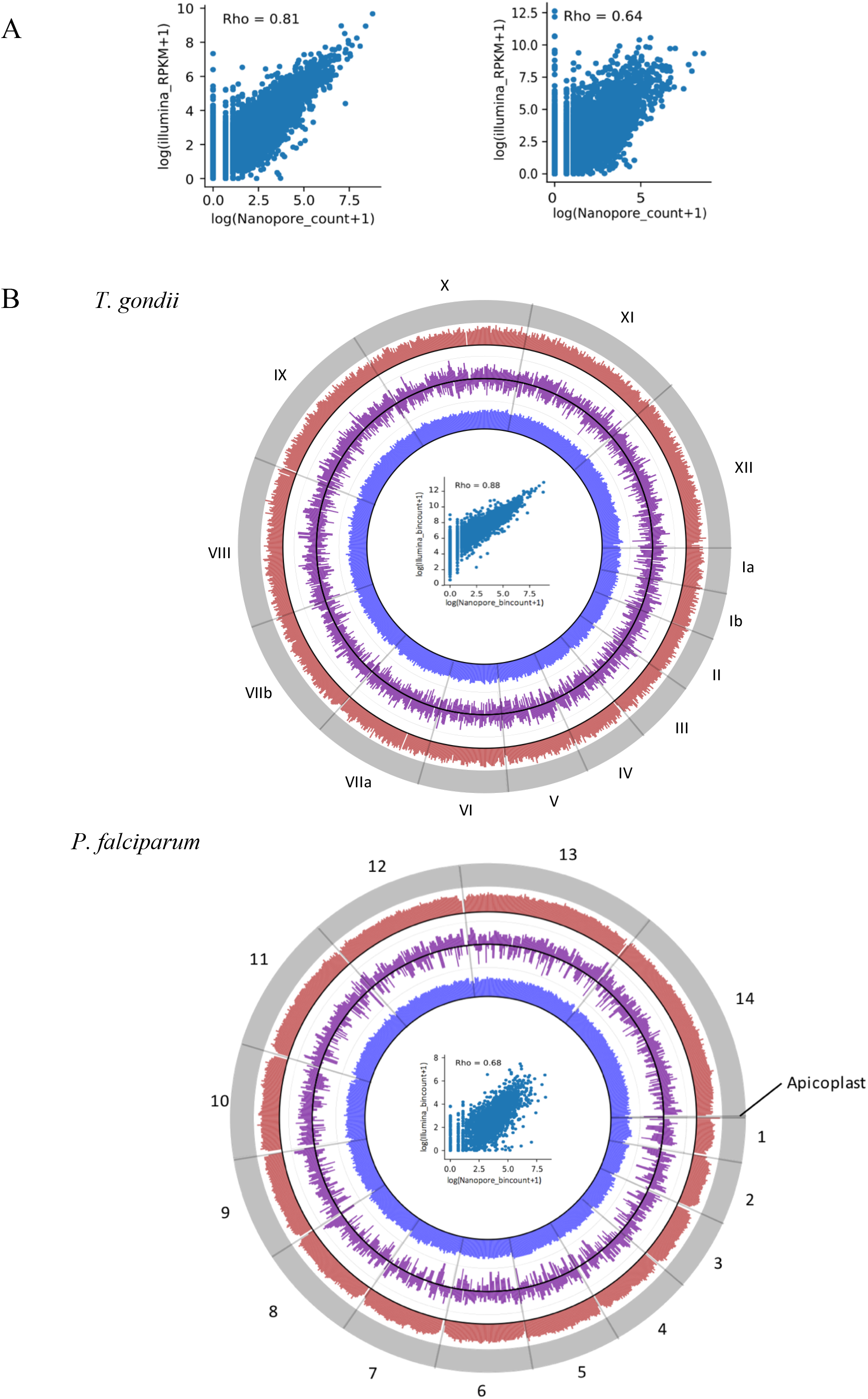
Comparisons between ONT direct RNA sequencing and Illumina datasets. (A) Correlation between trancriptome mapped read counts for *T. gondii* (left) and *P. falciparum* (right) as presented as a scatterplot. The Spearman correlation coefficient is shown. (B) Circos plots of genome mapped reads. Outer band (grey) represents the reference genome/chromosomes. The red and blue bands represent the genome coverage of ONT direct RNA and Illumina reads respectively. The purple band is the log2 fold change between the two datasets. The scatterplots within the circos plots show the correlation between the genome mapped read bin counts.

For both parasites, a higher number of gene transcripts were detected in the Illumina datasets than in the ONT data (supplementary table TS1). This is expected, given the greater sequencing depth that was obtained from the Illumina runs. The theoretical average fold coverage from our Illumina data is estimated to be 750 and 1000 times of the *T. gondii* and *P. falciparum* genes respectively (compared to 25-26 fold for the ONT data). Notably however, abundant non-mRNA species, particular ribosomal RNA, were detected in the Illumina datasets. This is absent in the ONT datasets, possibly because of differences in polyA RNA purification methodologies or biases due to PCR amplification. We further evaluated the ONT transcriptomes for genome coverage completeness as shown in Figure 2B. The read coverages from the ONT and Illumina reveal no significant bias towards a particular region of the nuclear genome.

### Intron retained transcripts are prevalent and generally non-productive

A major goal of mature full-length transcript sequencing is the identification of splicing isoforms. Alternative splicing can be broadly classed into four types: intron retention, alternative 3′ splice site selection, alternative 5’ splice site selection, and exon skipping [40]. Of these, intron retention is the least studied form of alternative splicing despite the numerous studies implicating the significance of the event [41–43]. This is in part due to the limitations of short read sequencing, but also the relatively long and low-complexity introns in metazoan genomes, which impose limitations on sequencing and assembly. For example, the intronic sequence in the human genome is several magnitudes longer than the length of the exonic sequence [44, 45]. In contrast, the compact genomes of *Plasmodium* and *Toxoplasma* both have gene models that have similar or longer exon lengths than introns [46, 47], so reads that span multiple entire introns are quite achievable for these organisms.

To monitor levels of intron retention, we identified junctions and reads that overlapped annotated intronic regions of a gene based on the annotated coordinates using FeatureCounts [48], and tallied the proportion of reads mapping to that intron to the total reads for the same gene. Proportion scores are represented using the metric *percent intron retention* (PIR). Based on this analysis, 17.65% of the mapped reads were considered to have intron overlaps for *T. gondii*, and 4.54% for *P. falciparum.* We further filtered out junctions without a minimum overlap of six bases to exclude artefacts generated by read errors, and excluded genes that were not supported by a minimum coverage of three reads. The distribution of PIR scores per gene (Figure 3A) reveals an overall skew towards low proportions of intron-overlapping reads. This is as expected given the propensity for a dominant canonical transcript [49]. However, we identify a strikingly high number of genes that retain a high level of intronic regions. Using a threshold of 10% PIR, we identified a total of 3229 genes for *T. gondii* and 978 genes for *P. falciparum* that have intronic reads within their transcripts (supplementary table TS2). Moreover, for around 29.82% (963/3229) of the *Toxoplasma* genes, and 19.63% (192/978) of *Plasmodium* genes, 50% or more of the reads retain at least one intron. Unusually, there are a considerable number of genes where none of the transcripts appears to have all of their annotated introns removed. We manually investigated these cases further and found major differences between the transcripts and gene model in most cases, suggesting that these highlight genes with incorrectly annotated structure. A couple of examples are outlined in supplementary figure S3. Most of these genes are annotated as hypothetical proteins, highlighting the potential of ONT sequencing to validate gene models.

**Figure 3.**
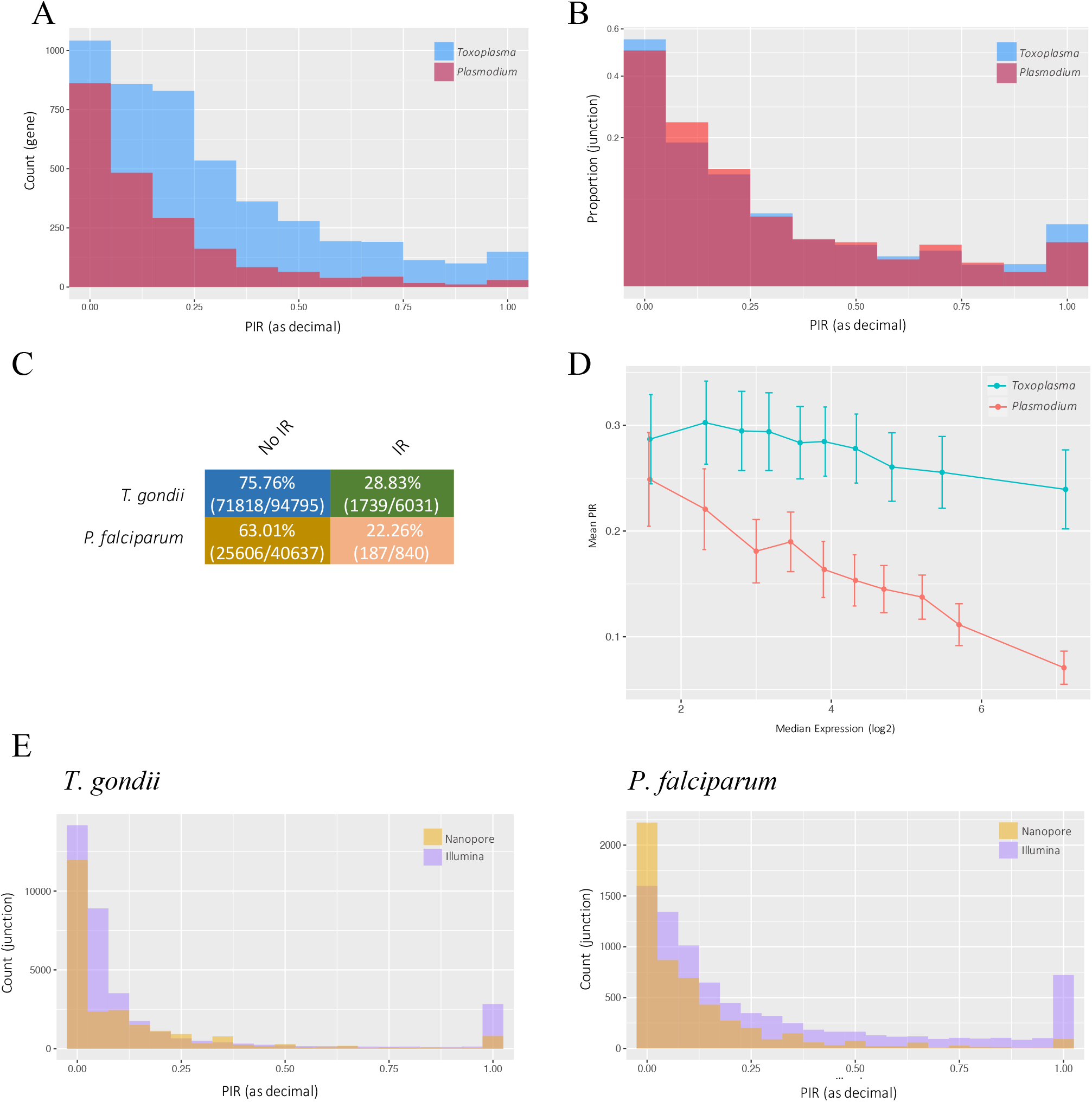
Analysis of intron retention using ONT direct RNA sequencing reads. Levels of intron retention are represented as Percent Intron Retention (PIR). (A) Distribution of intron retention levels over each gene as represented as total counts. (B) Distribution of intron retention levels over each junction as represented as the proportion of each bin count over the sum all intronic junctions. (C) Table of transcript productivity based on intron retention events as analysed using the FLAIR pipeline. Numbers in bracket are the transcript counts (D) Relationship between levels of intron retention and gene expression. Genes were classified into 10 bins of equal number based on expression. The median expression of the genes in each bin is used to represent expression for each bin. Error bars represent the 95% CI. (E) Comparisons of intron retention quantification between ONT and Illumina datasets.

The most extreme cases of conflict between the junctions we detect and canonical gene models often highlight potential annotation errors, but there are still a strikingly high number of genes where genuine introns are retained in a high (>50%) proportion of transcripts (*T. gondii*: 808 genes, *P. falciparum*: 162 genes). Additionally, the differences between the two parasites are striking. In many organisms, those transcripts with the most introns are those that are more likely to retain at least one or more introns [50]. This is partially supported in our analysis, where we observe a higher level of overall intron retention for *T. gondii* (which has 4.5 introns per gene on average) than *P. falciparum* (which has only 1.5 introns per gene on average). A possible explanation for this relationship is thus that both organisms have a similar level of intron retention for any given junction, and the higher average intron number in *T. gondii* genes results in more overall intron retention per gene. To examine this, we calculated PIR scores at the individual junction level, rather than per-gene level, and normalised the count of each PIR value to the proportion of total junctions within each organism. The analysis reveals that after correction for intron number there is virtually no difference in the distribution of intron retention levels between parasites (Fig. 3B). In other words, individual *T. gondii* junctions are no more likely to experience intron retention than *P. falciparum* junctions. We further tested whether intron number was the major predictor of intron retention at the gene level by looking at the correlation between the number of introns per gene and levels of intron retention. Interestingly, we only obtained poor or moderate positive correlations in all datasets (supplementary figure S4; Spearman’s rho = 0.28 for *T. gondii*, 0.48 for *P. falciparum*, 0.39 for pooled). This correlation does not significantly improve even when we restrict our analysis to higher (≥ 10 reads) expressed genes. This suggests that while intron number is associated with increased level of intron retention, it is not the main determinant of whether a gene retained at least one intron.

By taking advantage of the full-length reads made possible by ONT, we are able to predict the protein-coding productivity of the alternate transcripts. We performed productivity analysis on full-length intron-retained reads using the FLAIR [51] pipeline, which corrects and defines unproductive transcripts as transcripts with a termination codon that is 55 nucleotides or more upstream of the 3’-most splice junction. The rationale for this definition is based on previous evidence suggesting that only premature terminating transcripts following that 55-nucleotide rule mediate an effect on mRNA turnover [52]. This is a conservative estimate of productivity as it does not consider intron retention within the 3’-most splice junction. The Flair method identified over 70% of the intron-retained reads to be non-productive for either parasite (Fig 3C), suggesting that the high level of observed intron retention only rarely corresponds to alternative protein products, and that most intron-retaining transcripts may instead be targets for nonsense mediated decay. Intron retention is known to fine-tune protein expression through this pathway in mammalian systems [53]. A related prediction from other studies [54] is that the most highly expressed transcripts should have low levels of intron retention. In our analysis, we do observe a negative relationship between intron retention and gene expression levels (Fig. 3D). There is a relatively high variance for this, more so for genes with lower expression levels. This is likely due to the limitation in precision for the lower sequencing depths. For example, intron retention occurring in 10% of transcripts for a given gene will not be precisely measured for a gene for which only five reads are available. We circumvented this by classifying the transcripts into bins of equal read number based on expression and quantifying global intron retention levels within each bin. Again, we observe a negative relationship between intron retention and expression levels (supplementary figure S5). To investigate the functional significance of this, we further analysed the genes for Gene Ontology (GO) enrichment. Here, we only considered genes with a minimum coverage of 10 reads to increase precision. The analyses reveal the consistent enrichment of genes with functions associated with the ribosome when there are lower levels of intron retention across both parasites (supplementary table TS3). This association has been previously observed in other organisms [55], though its basis is unknown. We also tested whether intron retention correlated with essentiality based on previous functional genomic screens [56, 57], and found no significant relationships (supplementary figure S6).

To validate the identification of intron retention events, we looked at whether retained introns apparent in the ONT data were directly supported by Illumina RNAseq data. We normalised read counts by junction length and only considered intronic data that spanned the full junction. Based on the analysis, 77.88% for *T. gondii* and 87.37% for *P. falciparum* of the intronic junction reads flagged from the ONT datasets were supported by Illumina reads. However, we also noted that some alternative splicing events, particularly the lower frequency ones, failed to be captured by ONT sequencing compared to the Illumina dataset (Fig 3E). This is again likely due to the limitation in read depth in the ONT dataset. Based on our sequencing of polyA tailed material combined with previous kinetic studies [58, 59], we do not expect the intron retained transcripts to simply be unprocessed transcripts. To confirm this, we looked for evidence that each transcript had at least been partially processed. On average, 92.76% of multi-intron genes identified as having intron retention within their transcripts had at least 1 junction which was canonically spliced in all the transcripts, demonstrating that cannot be attributed to sequencing of pre-mRNA.

### Alternate junction splicing is often proximal and non-productive

Having previously identified intron-retained, read junctions using an annotated gene model approach, we used RSeQC to identify and quantify the other three classes of alternative splicing read junctions (exon skipping, 5’-, and 3’-splice site change) based on a similar methodology. Levels of alternative spliced junctions are calculated as the proportion of alternate junction over the total junction reads, and are represented using the metric percent spliced (PS). Here, we filtered out junctions unsupported by a minimum coverage of three reads. Using the same threshold as before (≥ 10%), we identified a total of 1138 genes for *T. gondii,* and 168 genes for *P. falciparum,* where one or more of their junctions exhibited alternative 5’/3’ splice site selection or exon skipping (supplementary table TS4_1 & 2). Remarkably, these aggregate numbers are lower than those we calculated for intron retention alone. Combining the datasets, intron retention accounts for 60-68% of alternatively spliced genes identified, alternate 5’ junction and 3’ junction splicing for 13-19% and 6-11% respectively, and exon skipping for less than 3% (Fig. 4). The rest of the junctions flagged in the analysis defy easy categorisation due to major mismatches between the RNAseq data and the annotated gene model. Subsets of genes were also found to have multiple alternative splicing type events within their transcripts as observed in supplementary figure S7, though there does not appear to be a particular functional trend.

**Figure 4.**
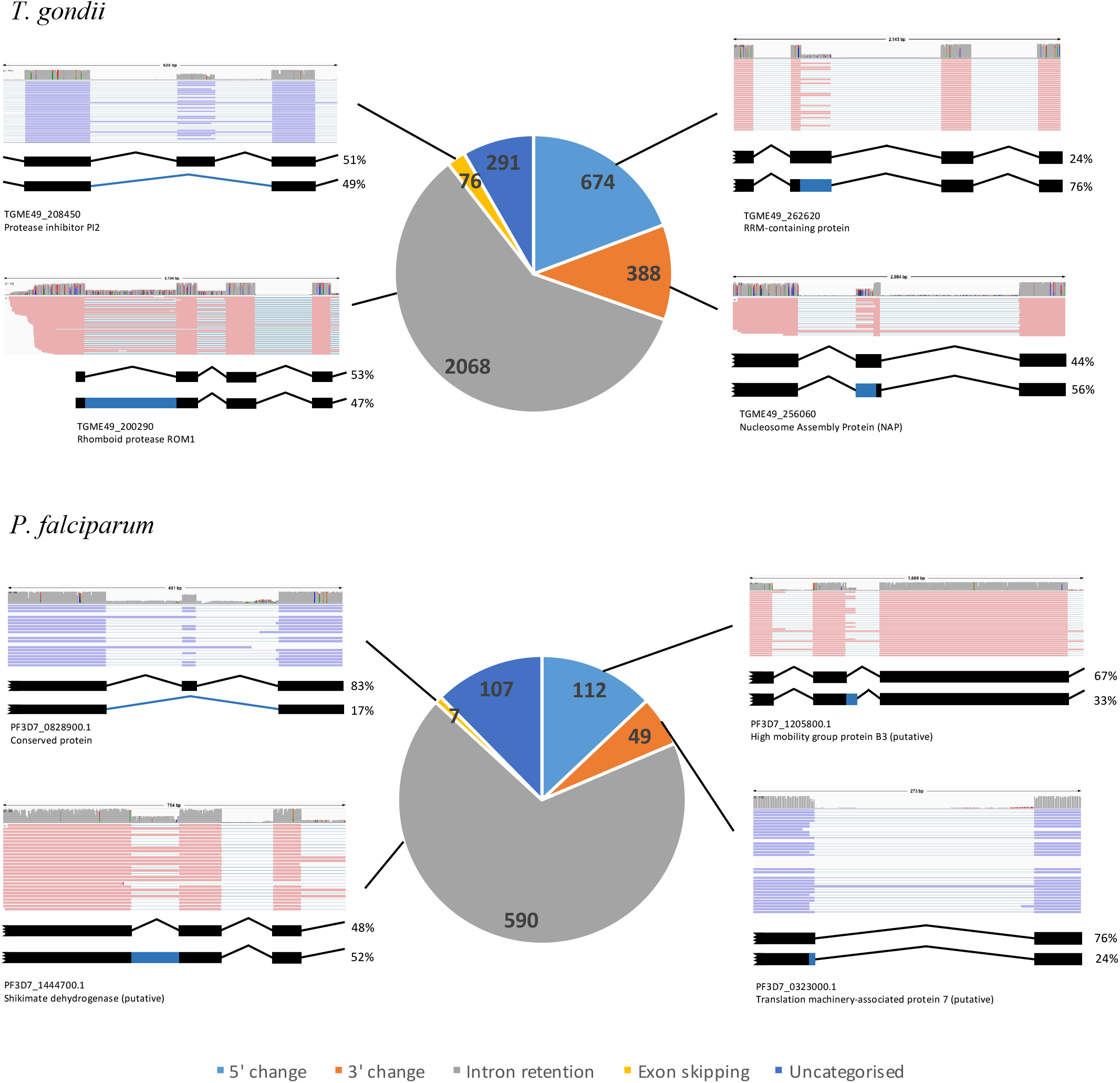
Summary analysis of alternative splicing from the ONT direct RNA sequencing datasets. Pie charts show the number, proportion and categorisation of genes with alternatively spliced transcripts equaling or exceeding 10% of its total transcript. An example of each event is presented. Red and blue represents sense and anti-sense transcripts respectively.

Having identified junctions subject to alternative splicing, we then quantified what proportion of the transcripts produced at those junctions represented the non-canonical isoform. For alternative splicing involving 5’ or 3’ changes and for intron retention, substantial proportions of isoforms were represented by the alternative transcript, but the median abundance remained below 50% (data shown in Fig. 5A). However, whilst exon skipping was a relatively rare event across the genome (Fig. 4), for those genes where exon skipping did occur, it represented a higher proportion (approximately two-thirds) of transcript isoforms for those genes than observed for the other forms of alternative splicing (Fig. 5A). We had previously published a list of genes with alternative splicing excluding intron retention in *T. gondii* (RH) using our in-house built computational tool– JunctionJuror [60]. A limitation of the tool over our current methodology is that the information about junction types is ultimately lost. Despite the differences in methodologies and parasite strain, we compared our list of genes with alternative splicing excluding intron with the previous list and found a moderately high degree of overlap (57.46 %) (supplementary table TS5). This intersect is double that is expected by chance alone and may represent genes with a steady state alternative splicing of higher confidence.

**Figure 5.**
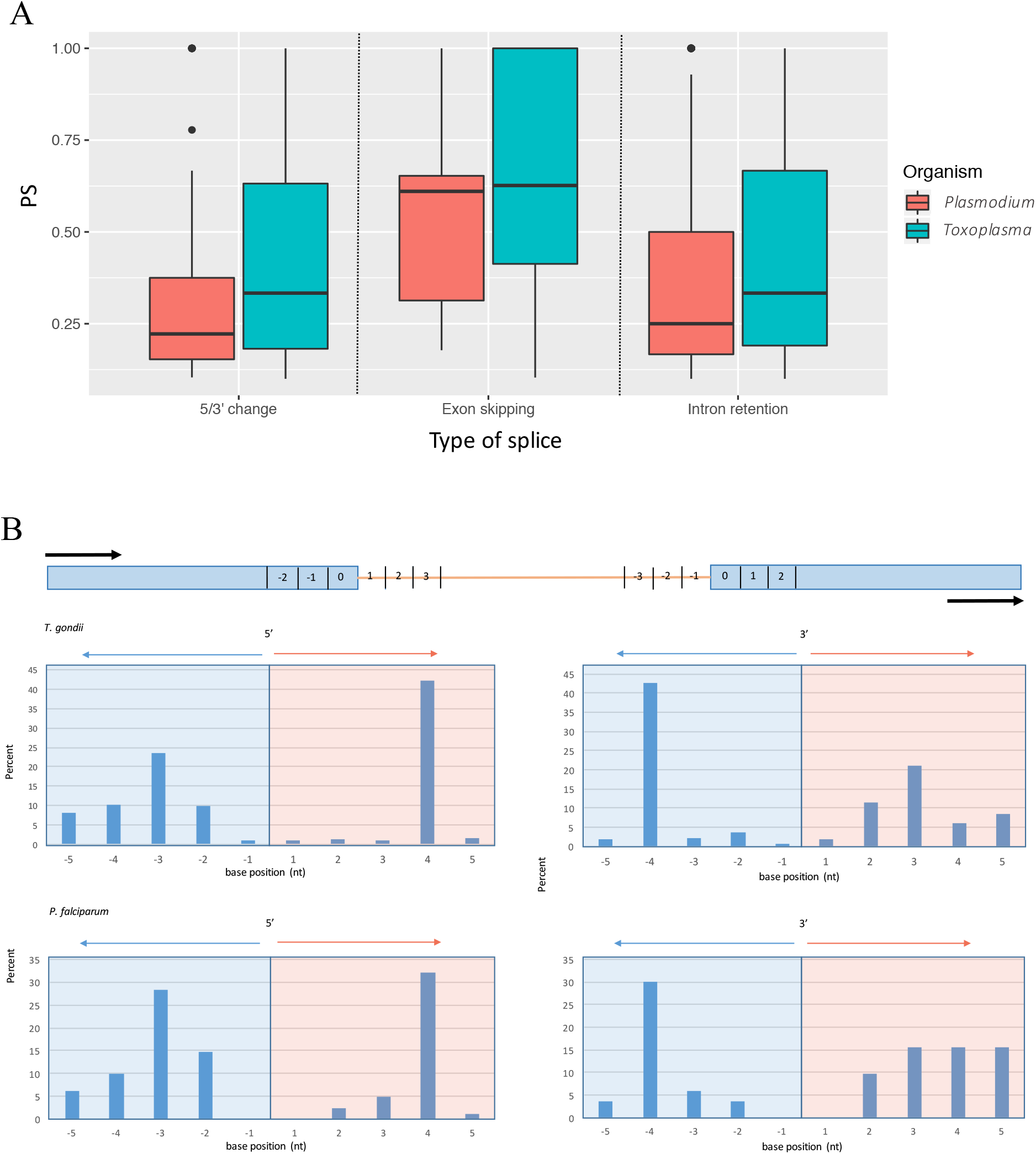
(A) Levels of each major alternative splice type as represented as Percent Splice (PS). Distribution of alternate 5’/3’ splice positions within 5 bases to the dominant splice site.

To further explore consequences of alternate 5’ or 3’ junction splicing events, we investigated the length distribution of the change in intron length. We found a surprisingly high proportion (~50%) of alternate 5’/3’ splicing to occur proximal (<6 bases) to the expected canonical site. We graphed the distribution of splicing change positions in Figure 5B and found a substantial spike of splicing changes occurring at the position four bases inside the canonical intron boundary. We tested these junction reads for productivity for a subset of 50 of these “near-miss” alternate splice events, and found all reads to prematurely terminate (unsurprisingly given the necessary frame-shift). This striking over-representation of isoforms departing from the canonical model by specifically four bases has been previously identified in metazoans [61]. Very small movements in splice site usage have been described as junction wobbling, and this has been proposed as minor splicing noise, or alternatively, as an additional mechanism of regulation through the NMD pathway [61], although the reason for the specific peak of AS four bases away from the canonical junction is unknown.

## Discussion

Despite the apparent significance of alternative splicing in metazoans, very little is known about the process in apicomplexans. A number of targeted experiments highlight single splicing events and their impact on parasite survival, but global splicing networks have been poorly described. Untargeted RNA-seq experiments have mainly focused on whole transcriptome assembly and/or gene expression. Those studies that do monitor alternative splicing reveal its occurrence in multiple genes and stages, but the extent of these events and their phenotypic significance remains unknown. The lack of a robust methodology in defining transcript isoforms from short read data is a particular challenge in dissecting whole-transcriptome splicing. In this study, we investigated whether ONT direct RNA sequencing could be used to explore the splicing landscape of two apicomplexan parasites- *T. gondii* and *P. falciparum.*

To our knowledge, ONT direct RNA sequencing has not been previously described in apicomplexans and so our first objective was to evaluate the capability of ONT sequencing in generating sequencing reads from these parasites. We successfully obtained high quality sequencing reads for both parasites that were comparable to that previously described in the literature for other organisms [62, 63]. In particular, we obtained read lengths that exceeded 1kb on average, many of them predicted to represent full-length or near full-length transcripts. Interestingly, although we obtained a higher number of sequencing reads for *P. falciparum,* the mapping of the reads was suboptimal compared to that of *T. gondii*. Repetitive DNA sequence motifs are characteristic of many large eukaryotic genomes and this has been known to complicate the mapping of reads that cannot be confidently assigned to these particular regions [64]. In theory, long read sequencing mitigates this problem because long enough reads should unambiguously match a unique site on the genome, irrespective of low complexity or repeat sequences. However, the genome of *P. falciparum* is particularly AT-rich (~82%) with numerous regions of extreme low complexity [65]. Thus, we may expect some reads, particularly the shorter ones, to fail the mapping parameters. Indeed, as indicated above, reads that fail to map to the genome tended to be shorter than reads that do. This is further exacerbated by the high error rate produced from ONT sequencing. Based on previous experiments, the per base error rate of direct RNA sequencing using ONT is 10-20% [51, 62]. In our dataset, we estimated the error rate to be around 20%. This may have further contributed to the poorer mapping, though we do not expect the high error rate to significantly impact our study because the main analysis is focused on splice connectivity, rather than base sequences. As has happened for ONT DNA sequencing, we are likely to see significant improvements in read and mapping accuracy of RNA sequences as improvements are made to the flow cell and base caller. A study carried out by Runtuwe and colleagues is an elegant example, where ONT DNA sequencing on targeted *P. falciparum* genes yielded a mapping percentage that improved from 57.86% to 92.46% with improved chemistry of the flow cell, and upgrades to the base-calling algorithms [66].

The quantification of gene expression is one important goal of RNA-seq. Traditional, short-read sequencing requires the generation and amplification of complementary DNA (cDNA) which can introduce artefacts and biases. Transcriptional amplification or repression is a commonly-overlooked bias [67], where the levels of global mRNA, rather than specific mRNA, may be variable between different samples. Thus, using a standard amount of total RNA, as is commonly done, can mask actual detection of specific mRNA levels, even after normalisation [67]. Direct RNA sequencing allows these caveats to be bypassed because a standard amount of isolated mRNA instead is used as the sequencing material. However, because there is no amplification step, direct RNA sequencing is limited by the amount of mRNA that can be practically obtained and used in the sequencing process. Without sufficient sequencing material, it can be difficult to achieve the high levels of sequencing depth that is needed to analyse gene expression [68]. In line with the literature, our analysis shows that the current protocol for ONT direct RNA sequencing is comparable to Illumina for quantifying gene expression in the organisms we analysed. It can be noted however that sequencing depth remains the main limitation of ONT sequencing in our study. The reduced throughput and sequencing depth from ONT sequencing compared to Illumina sequencing means that genes or transcript isoforms with low expression may not be captured.

Several previous analyses have reported differences in alternative splicing types and levels among different organisms [69–71]. Notably, the increase in intron number and its retention correlates strongly with multicellular complexity (as defined by numbers of distinct cell types) [72, 73]. In apicomplexans, the splicing machinery appears to be largely conserved but features of gene structure such as intron number, length and distribution can be highly variable [18]. In our study, the difference in intron number between *T. gondii* and *P. falciparum* is a relevant example. Despite the differences, we found that alternative splicing for any given junction occurs at similar rates between the two parasites. This further supports the notion that the parasites share similar splicing processes. Intron number of genes is predicted to be positively associated with alternative splicing events [50, 74]. This is not simply a stochastic effect but rather related to the general decrease in 5’ splice site strength with increasing intron number in many organisms [74]. As described above however, there is only a weak or moderately positive correlation between intron number and intron retention level of genes in the two parasites studied. Schmitz et al. [70] previously reported that other features such AT content and splice site entropy are important modulators of intron retention. This may also be the case in our study, given the observation that certain splice junctions are predisposed to retain their intron over others.

In addition to the difference in alternative splicing levels, there are differences in the composition of alternative splicing types between different organisms as reported by Kim and colleagues [71] and McGuire and colleagues [75]. For example, exon skipping is the predominant form of alternative splicing in metazoans, and intron retention the rarest [71]. Our analysis reveals that the opposite occurs in *T. gondii* and *P. falciparum*, with intron retention being the predominant event to occur, and exon skipping the rarest. This composition of alternative splicing type is similar to that observed for plants and fungi [71, 75, 76], though the reason is unclear. More recent studies find that intron retention has been previously under-detected in metazoans due to methodology limitations or confounding variables, but the high levels of exon skipping has been mostly undisputed [77]. Kim and colleagues [71] speculated that intron retention emerged as the earliest form of alternative splicing, before other mechanisms of complex splicing events evolved. There is some evidence for this, including the apparent shift towards increased exon skipping frequencies in early branching animals [77]. This is associated with the preservation of coding frames, suggesting a role of exon skipping in expanding proteome diversity [77]. In contrast to this, we found that the majority of the splicing events in *P. falciparum* and *T. gondii*, particularly intron retention, results in non-productive transcripts. Our results thus indicate that alternative splicing rarely contributes to generating diversity of protein sequence in these parasites, and may relate instead to transcriptomic complexity that impacts protein abundance. If that is true, a previous analysis that showed alternative splicing to be essential for *Plasmodium* stage differentiation [11] may possibly be explained by a requirement for modulation of abundance for specific proteins, rather than generation of protein sequences.

Consistent with our observation of alternative splicing playing a minor role in generating true proteome diversity in apicomplexans, many splicing events in other eukaryotes contribute little to the protein isoform repertoire. In particular, many transcripts contain premature termination codons (PTCs), at least in humans and yeast [78, 79]. Often, PTC transcripts are the result of the retention of intronic sequences that contain PTCs [80], but translational frameshifts from active splicing events such as alternative splice site selections have been similarly implicated [79, 81]. PTC transcripts are not normally translated but rather targeted for degradation through the nonsense-mediated decay (NMD) pathway [82, 83]. This is vital because the transcripts encode altered or truncated proteins which may exhibit deleterious activity [84]. Some studies postulate that the predicted alternative splice events are the result of either experimental or transcriptional noise [85], or that a substantial portion of such transcripts are contaminating pre-mRNA molecules, and so do not represent true alternative splicing [86, 87]. Nevertheless, many RNA-seq-based analyses operate on the assumption that PTC transcripts are biologically significant or relevant [88]. Congruously, studies focusing on mature mRNA isoforms in other organisms suggest that non-productive transcripts mediate an additional layer of post-transcriptional regulation, through downstream RNA processing changes such as mRNA turnover, export, and microRNA silencing [54, 89, 90]. Alternative splicing in apicomplexans may also play a role in these processes. Strikingly, unique PTC transcript signatures are associated with distinct cell lineages [42, 91, 92] in multicellular eukaryotes, which may be analogous to the essential role of alternative splicing observed in stage-differentiation observed in *Plasmodium* [11].

Non-productive transcripts are typically degraded though the NMD pathway, and this has been shown to regulate gene expression at the post-transcriptional level [93]. However, it is difficult to conclusively define the function of the non-productive transcripts without experimentally testing the proteomic fates of these transcripts. In metazoans, non-productive transcripts often highlight genes that were downregulated following a transition of cellular states [92]. Our study is consistent with this, given the observation that the number of non-productive transcripts generally decreased with increasing transcript number. This, in association with NMD, has been shown to be crucial to the maintenance and differentiation of many cell types [94, 95]. In contrast, in organisms such *Paramecium tetraurelia,* non-productive transcripts appear to be the result of splicing error rather than function [96]. Regardless, because gene expression as measured by transcript levels do not necessarily translate to protein expression levels, our findings have potential implications for the interpretation of RNA-seq studies in these parasites. Several studies have already demonstrated the poor correlation between protein and mRNA expression profiles in apicomplexans [7, 8, 97]. Our results highlight that for many genes, raw quantifications of transcript abundance will correlate poorly with the number of copies of productive isoforms, and provides one source of mismatch between transcriptional initiation and protein abundance.

Genome annotation is a crucial element of RNA-seq data analysis. For *T. gondii* and *P. falciparum*, the task is a widely accomplished manual effort from experts in the research community. Although genome annotation was not the main focus of the study, the ONT datasets are able to reveal the structure of full-length transcripts. This is crucial in validating gene models. Our data are viewable through the ToxoDB and PlasmoDB web resources [98], and raw data are available at the Sequence Read Archive (SRA) (https://www.ncbi.nlm.nih.gov/bioproject/PRJNA606986) which may aid the research community to further curate and validate the current annotations.

## Conclusion

In this study, we have performed the first direct transcriptomic analyses on *T. gondii* and *P. falciparum.* We show that ONT direct RNA sequencing enables the quantification of gene expression despite a reduced throughput. In combination with the increased requirement for starting material, this means that the cost and time per bp sequenced remains higher than that of second-generation sequencing platforms. Nevertheless, because ONT direct RNA sequencing enables the detection of full-length transcripts without amplification, the tool remains promising for resolving the limitations of second-generation sequencing.

We demonstrated that alternative splicing is widespread in the two parasites, particularly intron retention. ONT direct RNA sequencing enabled us to determine the productivity of these transcripts without complex computational methodologies, and we show that most the transcripts are premature terminating. This has implications for the quantification of gene expression, as it is highly unlikely for the wealth of transcript diversity that we identified to directly translate to protein isoforms.

## Methods

### Cell culture and RNA extraction

*Toxoplasma gondii* tachyzoites (Pru *Δku*80) were cultured on human foreskin fibroblasts in Dulbecco’s Modified Eagle medium (DME) supplemented with 1% v/v fetal calf serum and 1% v/v Glutamax. Freshly egressed tachyzoites were washed, filter purified (5um) and collected for RNA extraction. *P. falciparum* (3D7) were cultured in complete media consisting of human erythrocytes (O+, 2% haematocrit), RPMI-HEPES, 5% (w/v) Albumax and 3.6% (w/v) sodium bicarbonate. We collected mixed stage parasite, purified from host RBCs via lysis with 0.05% (w/v) saponin, for RNA extraction. To obtain the 500ng of mRNA recommended for the library preparation, we used TRI Reagent (Sigma) for extraction of total RNA followed by the Dynabeads mRNA Purification Kit for polyadenylated (poly-A) mRNA (Thermo Scientific) purification according to the manufacturer’s protocol. Purity and quantification of mRNA were determined via NanoDrop (Thermo Scientific) and a Qubit RNA HS Assay kit (Thermo Scientific).

### Library prep and Nanopore sequencing

Libraries for the direct RNA sequencing were generated using the recommended protocol for the SQK-RNA001 kit (Oxford Nanopore Technologies). We loaded and sequenced the libraries on MinION R9.4 flow cells (Oxford Nanopore Technologies) for 48 hours. Base calling was performed concurrent with sequencing using Albacore (v 2.0), which was integrated within the MinION software (MinKNOW, v1.10.23). Only “pass” reads were used for subsequent analyses.

### Mapping and qualitative analysis

ONT sequencing data was first checked for quality with FastQC (v.0.11.7) [99]. We then utilised Minimap2 (v. 2.1) [38] to map raw reads to the parasite genome and transcriptome from ToxoDB and PlasmoDB (r. 39), using the recommended preset commands. Intron length thresholds were set at 5000 and 1500 bases for *T. gondii* and *P. falciparum* respectively. Previously published Illumina datasets (SRR350746, ERR174301, ERR185969, ERR185970, ERR185971) were mapped using HISAT2 [100] using the preset commands. We checked for mapping quality with Samtools (v.1.7) [101], Picard (v.2.18.2) [102] and AlignQC (v.1.2) [103]. Further qualitative or quantitative analyses and graphical elements were done using the command-line interface and RStudio. We verified and illustrated subsets of mapped reads via IGV [104].

We correlated the ONT sequencing data with the Illumina datasets as previously described [105] using the wub package (v.0.2) [106]. The genome coverage of sequencing datasets were generated using bedtools genome coverage (v2.27) [107] and visualised via J-Circos (v1) [108]. Log2-fold ratios were calculated using DeepTools bamCompare (v.2.5.1) [109].

### Alternative splicing analysis

We applied two approaches to analysing alternative splicing. We first identified intron retained junctions and transcripts using featureCounts (v.1.6.2) [48] on the genome mapped reads. featureCounts matches features specified in an annotation file (gff) to mapped reads. The annotation files used in the analyses were obtained from ToxoDB and PlasmoDB (r. 39), and preprocessed via ToolShed (v.1.0) [110] to specifically extract intron coordinates and gene IDs. We set a minimum threshold requiring mapping to at least six-bases of the intron feature, and a minimum threshold of three reads mapping to the junction/transcript to be considered for further analysis. Percent Intron Retention (PIR) scores were calculated as the proportion of alternative splicing events to the sum of reads for each junction/gene. Productivity of full length transcripts was analysed using the Flair pipeline [51] using default parameters.

For the second approach, we used the junction_annotation.py script of RSeQC (v.2.6.4) [111] to identify novel or partial-novel junctions from the genome mapped reads based on the unmodified annotation file. Again, we filtered out junctions that had fewer than three supporting reads. The junctions were summarised into a table based on coordinate matching to the 5’ and/or 3’ of the expected canonical junction. We identified alternative 5’/3’ splicing and exon skipping based on the coordinates and strandedness of junctions identified by RSeQC that were either consistent with or conflicted with the annotated junctions. We manually validated the data, matched junctions to available gene IDs, and again calculated the proportion of alternative splicing events to the sum of reads for each junction. Using the final dataset, we re-curated the list of intron-retained junctions to exclude for alternate 5’/3’ splice changes. Proportional Venn diagrams were drawn using BioVenn [112].

Gene set enrichment analyses were carried out by ranking the genes based on their alternative splicing levels and using the first and third quartile of the ranked list as input for GO enrichment analysis via ToxoDB/PlasmoDB based on curated and computed assigned associations. We required the adjusted p-value to be smaller than 0.05 and FDR q-value of less than 0.25. This approach was validated using GSEA via WebGestalt [113].

## Supplementary files

Supplementary table TS1. An Excel spreadsheet quantifying the expression of genes from Illumina and ONT data.

Supplementary table TS2. An Excel spreadsheet quantifying the levels of intron retention at the gene level.

Supplementary table TS3. An Excel spreadsheet summarising the GO enrichment results for genes with high and low levels of intron retention.

Supplementary table TS4_1. An Excel spreadsheet quantifying categories of alternative splicing at the junction level for *T. gondii.*

Supplementary table TS4_2. An Excel spreadsheet quantifying categories of alternative splicing at the junction level for *P. falciparum.*

Supplementary table TS5. An Excel spreadsheet listing previously and currently identified genes with alternative splicing excluding intron retention for *T. gondii.*

---

**Supplementary Figure S1.**
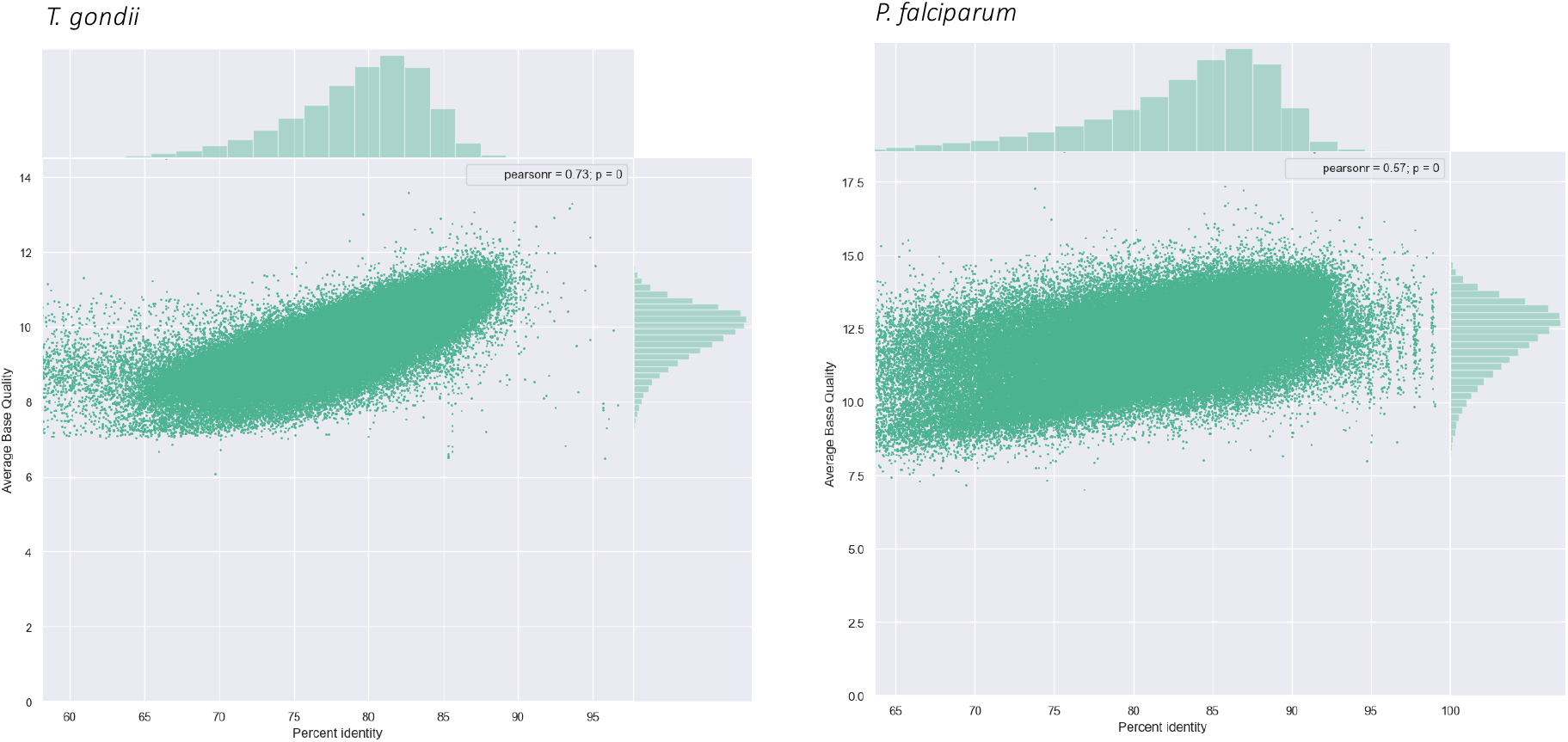
Scatterplot shows the correlation between transcript identity (%) and average quality of the reads.

**Supplementary Figure S2.**
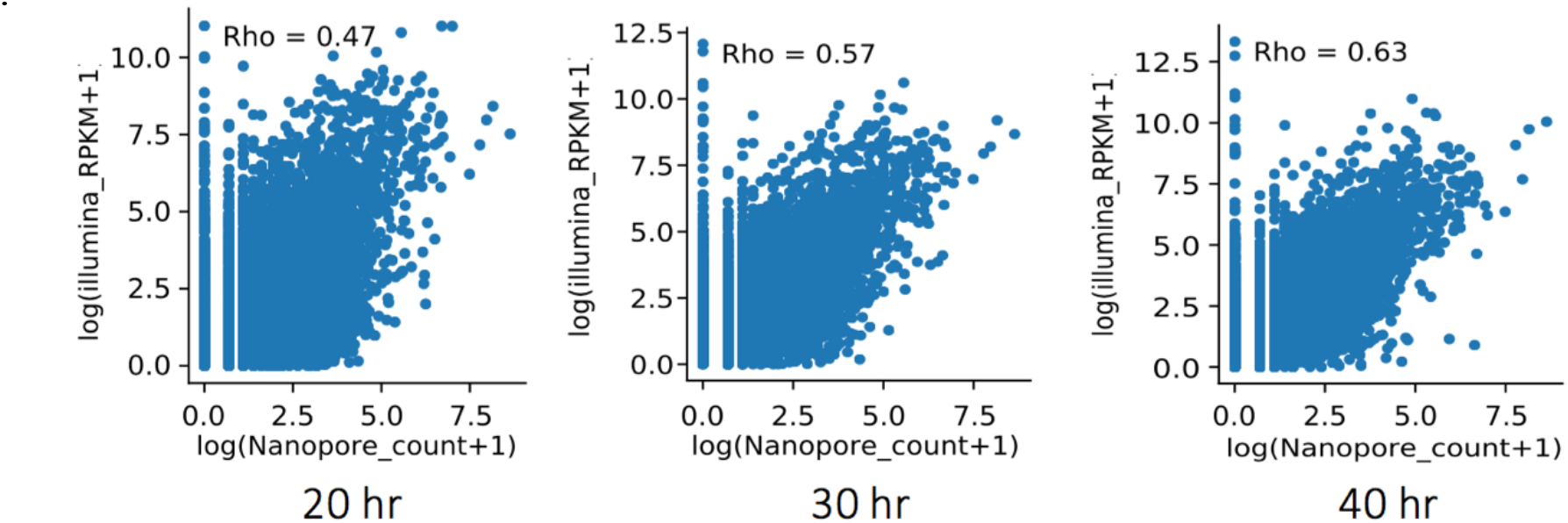
Scatterplot shows the correlation between trancriptome mapped read counts for ONT direct RNA sequencing from mixed stage parasites and Illumina datasets from three developmental time points of *P. falciparum*. The Spearman correlation coefficient is shown.

**Supplementary Figure S3.**
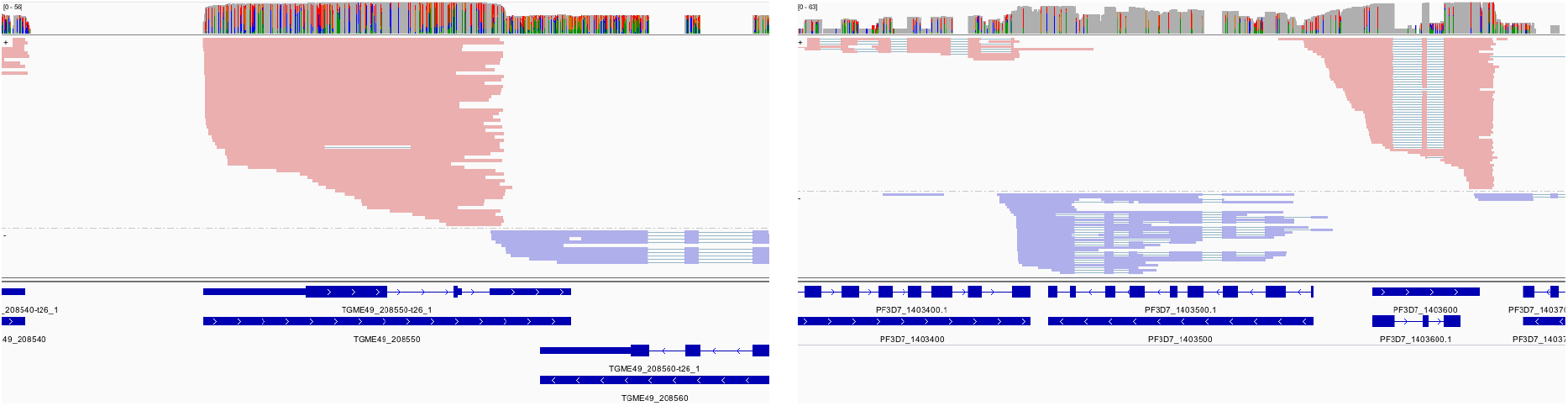
IGV snapshots of genes which were flagged as having full intron retention and do not appear to conform with the gene model. Left: *T. gondii*; Right: *P. falciparum.*

**Supplementary Figure S4.**
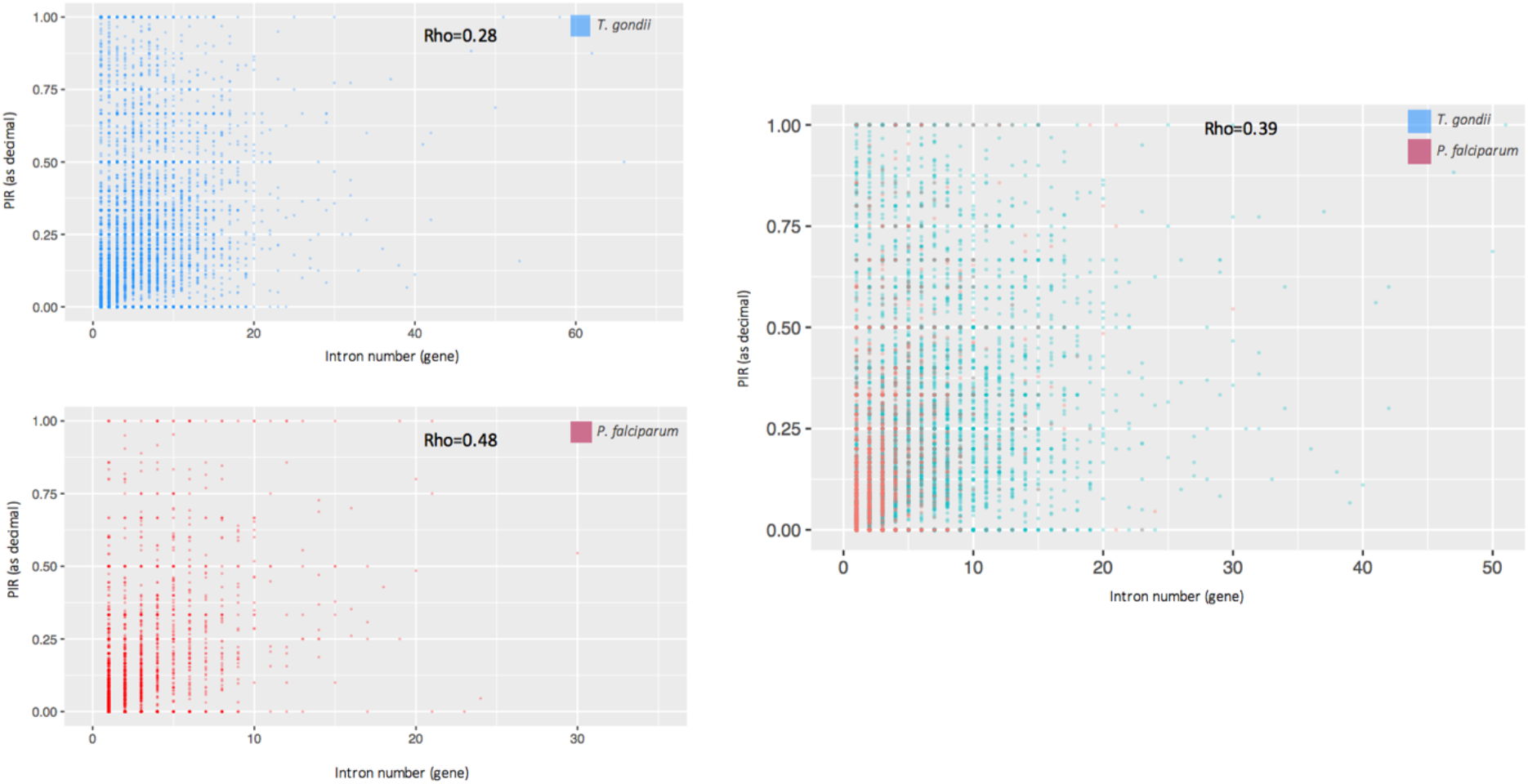
Scatterplots show the correlation between intron retention levels and intron number per gene. The Spearman correlation coefficient is shown.

**Supplementary Figure S5.**
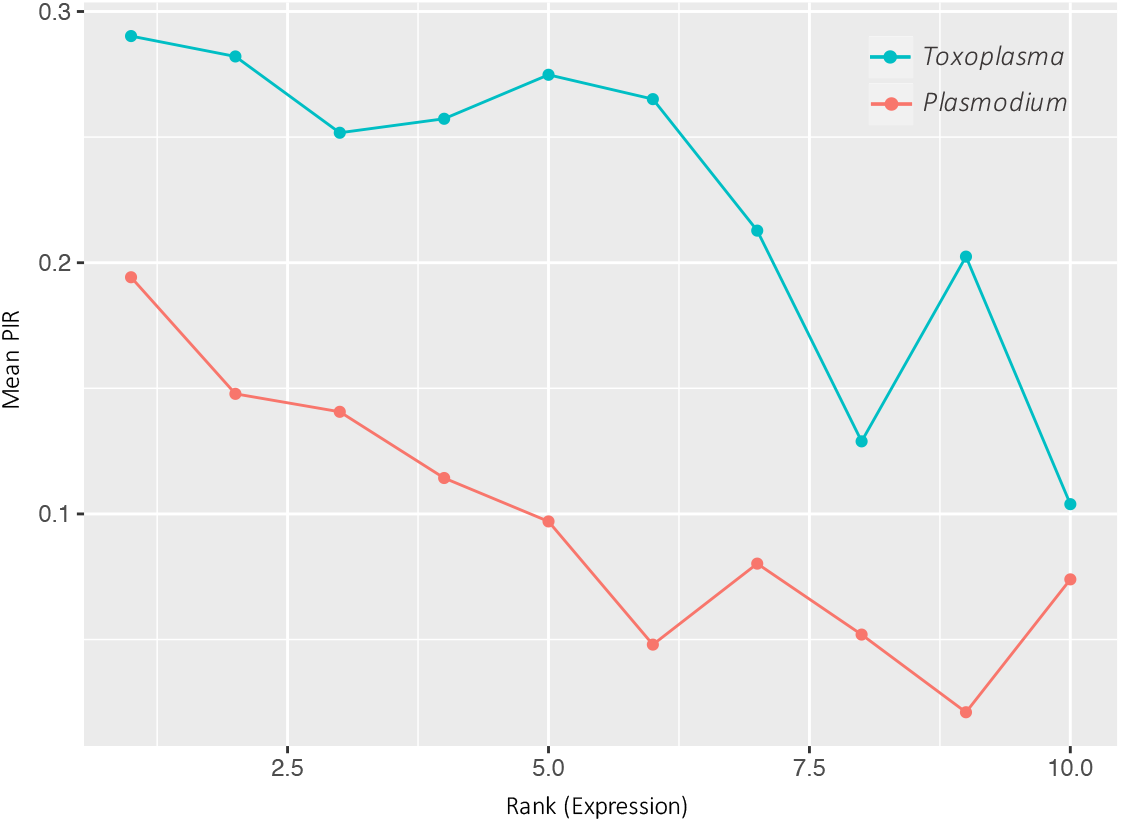
Relationship between levels of intron retention and gene expression. Introns were classified into 10 bins of equal read number based on expression (1 = lowest, 10 = highest). Levels of intron retention were calculated as the proportion of intron retained reads over all reads from each bin.

**Supplementary Figure S6.**
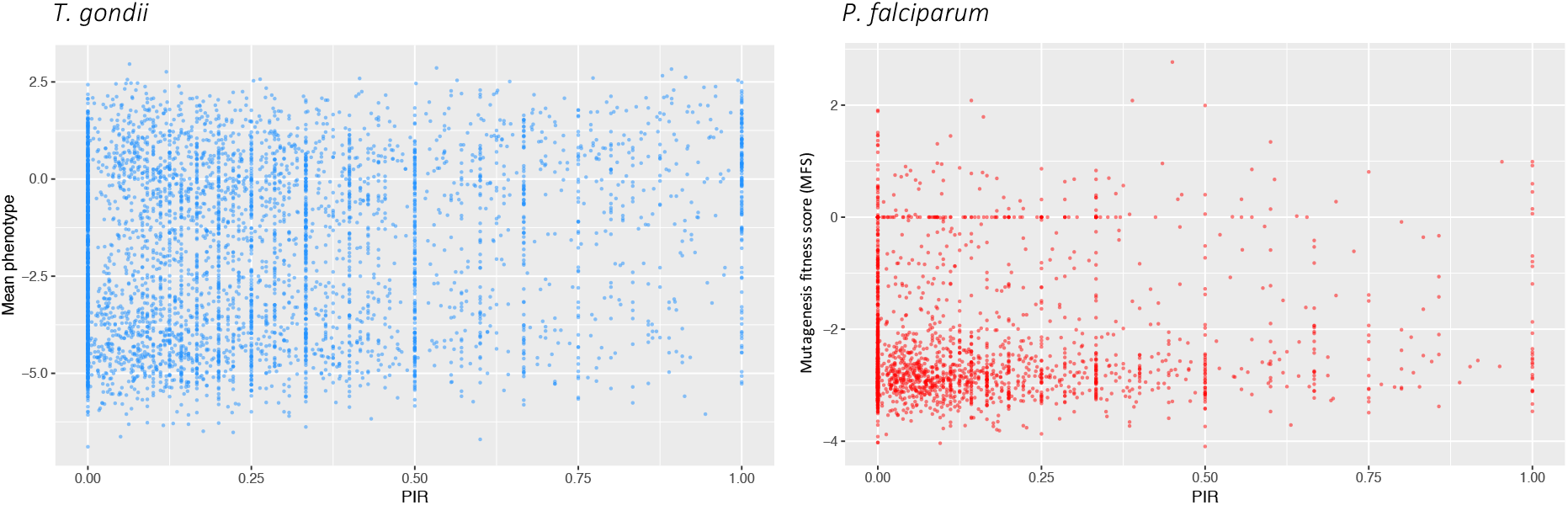
Scatterplots show the relationship between intron retention levels and gene essentiality. Essentiality is represented as mean phenotype and mutagenesis fitness score for *T. gondii* and *P. falciparum* respectively

**Supplementary Figure S7.**
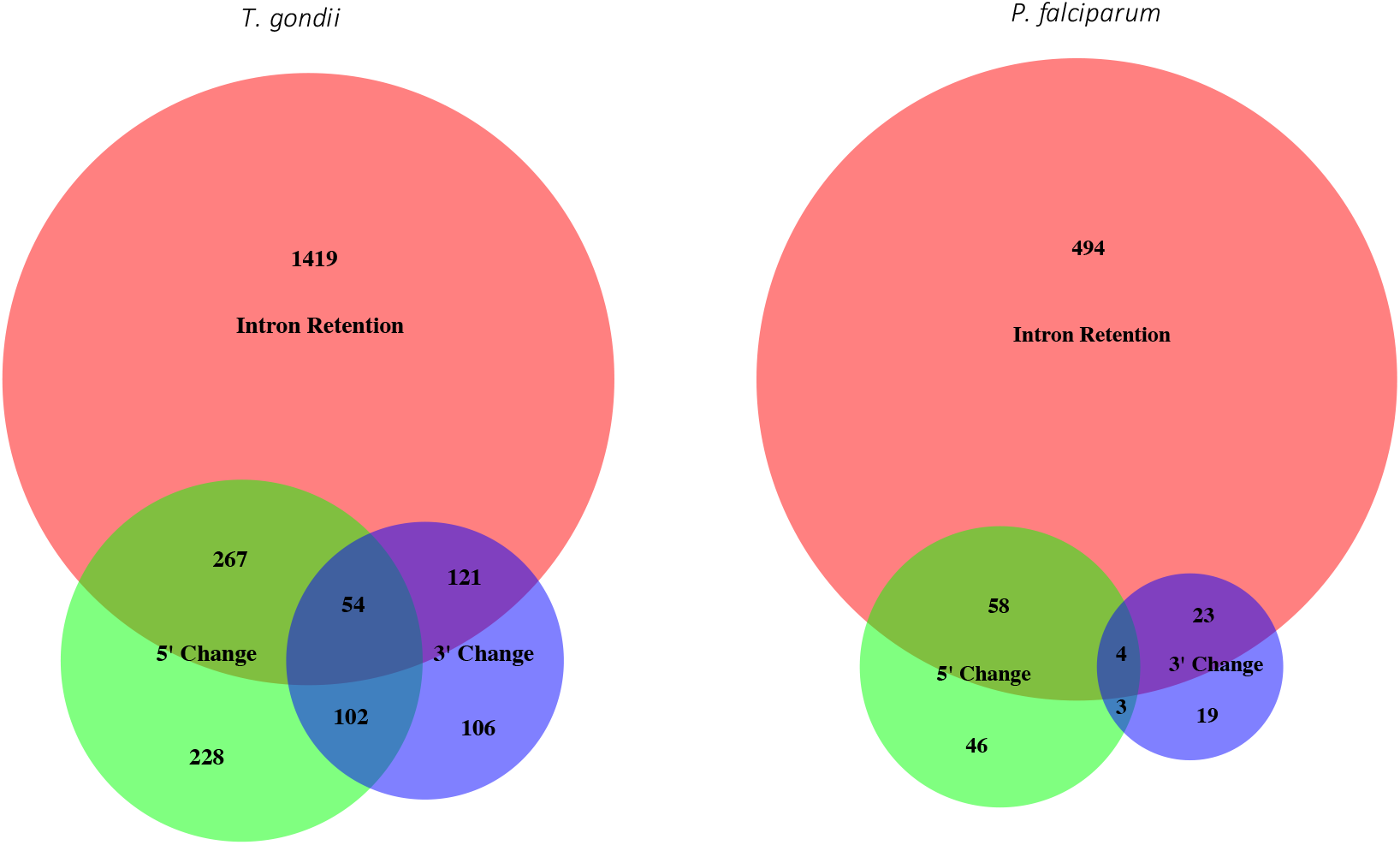
Venn diagrams display the number of genes with exclusive or overlapping intron retention, alternative 5’ splicing and alternative 3’ splicing events.

## Notes

### Competing Interest Statement

The authors have declared no competing interest.

